# Extensive remodeling of the ubiquitination landscape during aging in *C. elegans*

**DOI:** 10.1101/2023.12.07.570635

**Authors:** Seda Koyuncu, David Vilchez

## Abstract

In previous work, we investigated ubiquitination changes during aging in *C. elegans*. We identified 2,163 peptides undergoing age-related ubiquitination changes in wild-type animals, corresponding to 1,050 proteins. While many lysine sites had increased ubiquitination with age, a larger number exhibited decreased ubiquitination. Longevity pathways, such as reduced insulin signaling and dietary restriction, prevented ubiquitination changes. Treatment with a broad-spectrum inhibitor of deubiquitinating enzymes (DUBs) or knockdown of specific DUBs ameliorated ubiquitination loss in old worms. Moreover, we identified proteins that accumulate with aging due to reduced ubiquitination and subsequent proteasomal degradation. Our conclusions were supported by multiple approaches, including ubiquitin and total proteomics, western blot, and ubiquitin-less mutations. Concerns were raised by a laboratory regarding the possibility of our protocol omitting insoluble proteins due to a centrifugation step after protein extraction and solubilization. To address these concerns, we have focused on proteins previously reported to become insoluble with aging in *C. elegans*. Our proteomics experiments successfully detected and quantified all these proteins in old worms. In many cases, the proteins that become insoluble with aging did not change or even increased at the total levels. However, they often exhibited ubiquitination changes, primarily a loss of ubiquitination. Independent work combining analysis of conformational changes with our datasets demonstrated that 92% of the age-dependent metastable proteins exhibit differential ubiquitination during aging. In addition, we performed experiments to confirm that the buffers used for proteomics and western blot efficiently solubilize most of the proteome. Importantly, the analysis of total homogenates combining clear lysates and the remaining debris after protein extraction also revealed decreased ubiquitination during aging, whereas DUB inhibitor treatment (4 h) in old worms restored ubiquitination levels. These data further support our previous conclusions regarding extensive ubiquitination changes during aging in *C. elegans*.

## Introduction

We are writing in response to the recent preprint by Daigle et al., available on bioRxiv^1^. In *C. elegans*, different proteins become insoluble during aging, leading to their aggregation^2-4^. The presence of ubiquitin has been detected in aggregates of mutant proteins linked to neurodegenerative diseases such as Alzheimer’s, Huntington’s and amyotrophic lateral sclerosis^5^. Daigle et al. claim that in Koyuncu et al.^6^ we largely omitted polyubiquitinated proteins that accumulate as an insoluble fraction. To our knowledge, there are no data indicating that proteins aggregating with age in *C. elegans* are generally polyubiquitinated^2-4^. In fact, the proteasome, which is the primary eukaryotic machinery for recognizing and degrading ubiquitinated proteins, exhibits increased activity with age in *C. elegans*^4^.

In our original publication, we discovered that aging induces ubiquitination changes across the proteome in *C. elegans*^6^. While many peptides undergo increased ubiquitination with age, a greater number exhibit decreased ubiquitination. Importantly, longevity pathways such as reduced insulin signaling and dietary restriction prevent these ubiquitination changes. Moreover, the treatment with a broad-spectrum inhibitor of deubiquitinating enzymes (DUBs) rescues loss of ubiquitination in old worms. Likewise, knockdown of specific upregulated DUBs also ameliorates loss of ubiquitination during aging. We identified a subset of proteins that accumulate during aging owing to loss of ubiquitination and subsequent degradation by the proteasome. Our data demonstrated that the accumulation of these proteins dysregulates cell function and contributes to organismal aging. Our conclusions were based on the combination of multiple approaches, including ubiquitin and total proteomics, western blot, and ubiquitin-less mutations. Importantly, a recent study has shown an extensive remodeling of the ubiquitination landscape in the mouse and killifish brain during aging^7^. This work extends our previous findings in worms to the vertebrate brain, supporting that remodeling of protein ubiquitination is a major signature of aging across species^7^.

Since we performed a centrifugation step to remove debris after protein extraction and solubilization, Daigle et al. argue that our study omitted insoluble proteins^1^. Here we show that proteins that become insoluble with age were detected and quantified in our experiments. In fact, we found that the total levels of many of these proteins remain similar or even increase during aging, contradicting the claim that insoluble proteins were largely omitted from our analysis. Moreover, we have performed additional experiments to further validate our interpretation and conclusions regarding extensive remodeling of the ubiquitinated proteome during aging. Indeed, we confirmed that our lysis protocols efficiently solubilize most of the proteome. In addition, we observed a similar loss of ubiquitination when we analyzed clear lysates or total lysates including debris after protein solubilization. We now update the community on these new results so they can evaluate our conclusions.

### Our proteomics datasets include aggregating proteins during aging

In their manuscript, Daigle et al used a harsh lysis procedure, involving a 5% SDS buffer followed by heating (95 °C, 5 min) and sonication. After centrifugation, the supernatant was collected and the remaining pellet was solubilized in an equal volume of buffer using the same procedure (i.e., 5% SDS buffer, heating and sonication). Then, the authors combined equal volumes of supernatant and pellet to analyze the total levels of ubiquitinated proteins by western blot. The authors also tested what they referred to as “mild homogenization,” resembling our buffer and procedure for western blot. In this case, they used a lysis buffer containing 1% NP-40 and 0.25% sodium deoxycholate as detergents. NP-40 is a relatively mild, nonionic detergent effective at solubilizing membrane proteins and isolating cytoplasmic proteins. Sodium deoxycholate is a denaturing ionic detergent used to completely extract and solubilize proteins from the nucleus and also insoluble fractions or aggregates. The percentage of sodium deoxycholate (0.25%) is the same as that used in our manuscript, within the typical range of 0.1% to 0.5% for lysing insoluble proteins. After lysis and centrifugation, Daigle et al. separated the supernatant and the remaining pellet was solubilized in an equal volume of 5% SDS lysis buffer, followed by heating and sonication. For western blot analysis, the authors loaded a mixture containing equal volume of supernatant and pellet.

As reported in our manuscript^6^, we only used NP-40 detergent for immunoprecipitation of proteins containing Lys48- and Lys63-linked polyubiquitin chains. For western blot of global ubiquitination levels, we used RIPA buffer containing 0.25% sodium deoxycholate and 1% Triton X-100 as detergents. We inadvertently omitted information regarding the use of Triton X-100 in the Methods section, and we have submitted an Author Correction to the journal. We sincerely apologize for any inconvenience this oversight may have caused. It is worth noting that since Triton X-100 and NP-40 can yield similar results, we believe this correction does not impact the concerns raised by Daigle et al., as they used NP-40 in their experiments.

For western blot analysis, Daigle et al. used the same antibody against ubiquitin as employed in our manuscript (Sigma, 05-944, clone P4D1-A11). Although the authors assert that they “unequivocally observe that polyubiquitinated protein levels do not decrease with age in *C. elegans*”, three out of the four blots presented in their manuscript indicate less ubiquitination in day 15-adult worms when compared with day-5 adults^1^. Nevertheless, we respectfully think it is difficult to reach conclusions from the western blot images presented in the preprint. For instance, the authors did not state how many independent experiments they performed for each buffer. They used actin as a loading control, but the amounts of actin across samples are different, making it difficult to interpret the results. The authors indicate that they collected approximately 500 worms per time point for each solubilization method. This is an important difference compared to our protocol, as we normalized the samples by total protein whereas Daigle et al. normalized the samples per number of worms. Since the goal of the study is to analyze ubiquitination levels in global protein extracts, it is relevant to load equal amounts of total protein.

Nevertheless, Daigle et al. assert that their western blots invalidate our conclusion regarding changes in the ubiquitinated proteome during aging in *C. elegans*. We respectfully disagree with this statement, based on the comprehensive data integrating different approaches presented in our original publication and new experiments performed in response to Daigle et al. First, it is important to highlight that the primary evidence supporting a remodeling of the ubiquitinated proteome during aging stems from our ubiquitin proteomics data. In these experiments, we used an antibody that recognizes di-glycine moieties linked by an isopeptide bond to lysine sites of proteins. These epitopes, remnants of ubiquitination following tryptic digestion, can be isolated biochemically and analyzed using mass spectrometry. In contrast to western blot experiments, ubiquitin proteomics enables the definition of ubiquitination changes at the resolution of individual proteins in a systematic manner. This technique has been used in multiple publications to provide site-specific information and quantification of ubiquitin modifications across the proteome^8-10^.

For proteomics experiments, we used a lysis buffer containing 10 M urea^6^. Urea is an effective denaturant to solubilize proteins, preventing protein precipitation and aggregation^11,12^. In our original study, we applied ubiquitin proteomics to compare wild-type worms at different ages (day 1, 5, 10, and 15) as well as age-matched genetic models of dietary restriction (*eat-2(ad1116)*) and reduced insulin signaling (*daf-2(e1370)*). With high reproducibility between replicates, we identified and quantified ubiquitination sites for 3,373 peptides that correspond to 1,485 proteins^6^. We found multiple changes in the ubiquitinated proteome of wild-type worms during aging. A detail analysis of changes in individual proteins provides further evidence of alterations in the ubiquitinated proteome during aging. Here, we will focus on the comparison between day-5 (young) and day-15 (old) wild-type worms (please see Supplementary Table 4 of our original publication^6^). In aged worms, we identified a total of 2,163 peptides that exhibit significant changes in ubiquitination with age, demonstrating a remodeling of the ubiquitinated proteome. More specifically, 350 Ub-peptides were upregulated, while 1,813 Ub-peptides were downregulated, supporting our conclusion that aging is particularly associated with a loss of ubiquitination in *C. elegans*. However, reduced insulin signaling and dietary restriction prevented most of these changes.

The 2,163 peptides undergoing age-related ubiquitination changes in wild-type animals correspond to 1,050 distinct proteins. In multiple proteins, we observed lysine sites that displayed reduced ubiquitination, while other sites became more ubiquitinated or remained similar within the same protein. In **Table 1**, we have presented examples of proteins undergoing ubiquitination changes in different directions depending on the lysine site, including chaperones and components of myosin filaments. Given that many proteins exhibit ubiquitination changes in different directions along with lysine sites that remain similar, our proteomics data demonstrate changes in specific sites during aging.

**Table 1.** Examples of proteins that undergo ubiquitination changes in multiple lysine sites during aging in wild-type *C. elegans*. Number of total, upregulated, downregulated and non-significantly changed Ub-modified lysine sites in different proteins when comparing day-15 and day-5 adult wild-type worms (n = 4; two-sided t-test, false discovery rate (FDR) < 0.05 was considered significant). The table also indicates whether the total levels of the corresponding protein change during aging. The data was obtained from the proteomics experiments and analysis presented in Koyuncu et al (2021)^6^.

| Protein | Total number of Ub-sites | Upregulated Ub-sites | Downregulated Ub-sites | Ub-sites with no changes | Total protein levels |
| --- | --- | --- | --- | --- | --- |
| HSP-1 | 25 | 0 | 17 | 8 | Decreased |
| HSP-25 | 5 | 1 | 3 | 1 | No change |
| HSP-43 | 4 | 1 | 1 | 2 | Increased |
| SQST-1 | 18 | 1 | 8 | 9 | No change |
| MYO-2 | 13 | 1 | 11 | 1 | No change |
| MYO-3 | 15 | 3 | 6 | 6 | Increased |
| MYO-5 | 10 | 2 | 3 | 5 | No change |
| UNC-15 | 34 | 9 | 16 | 9 | No change |
| UNC-54 | 35 | 4 | 31 | 0 | Decreased |
| ICD-1 | 8 | 1 | 7 | 0 | Decreased |
| LEA-1 | 102 | 6 | 78 | 18 | Decreased |
| PUD-2.1 | 11 | 0 | 8 | 3 | Decreased |
| VIT-6 | 31 | 1 | 19 | 11 | No change |

In parallel, we quantified the total amounts of individual proteins for comparison with Ub-peptide levels^6^. Among the 350 upregulated Ub-peptides, 123 peptides correlated with an increase in the total levels of the respective protein, whereas 180 peptides corresponded to proteins that remain similar in abundance with age. Likewise, over half of the 1,813 downregulated Ub-peptides corresponded to proteins that maintain similar total levels during aging^6^. The intermediate filament IFB-2 is a protein that we found to aggregate with aging, while its lysine sites become less ubiquitinated. Despite that IFB-2 aggregated during aging, we observed increased levels of IFB-2 protein in the lysates analyzed by proteomics and western blot^6^. These data further demonstrate that we are monitoring changes at the global proteome level regardless of solubility.

To further address the concern from Daigle et al. that our protein extraction overlooked insoluble proteins, we have now specifically focused on proteins known to aggregate during aging (**Table 2**). For instance, IFC-2 is another intermediate filament that we found to accumulate into insoluble aggregates during aging in our original publication^6^. Since IFC-2 was detected in our proteomics experiments, and its total protein levels remained unchanged with age (**Table 2**), these data contradict the claim that we omitted insoluble proteins in our experiments.

**Table 2.** Proteins that are known to aggregate during aging in *C. elegans* were detected in our ubiquitin and total proteomics experiments. The table shows a list of bona-fide examples of proteins that aggregate during aging in *C. elegans* (David et al. and Reis-Rodrigues et al.^2,3^). Importantly, we could quantify all these insoluble proteins even in day 15-adult worms using our lysis procedure. Most of these proteins present changes in ubiquitination when comparing day-15 and day-5 adult wild-type worms, but we also observed ubiquitination sites that remain similar (n = 4; two-sided t-test, false discovery rate (FDR) < 0.05 was considered significant). The total levels of many of the aggregation-prone proteins were not changed or even increased during aging (n = 4; two-sided t-test, false discovery rate (FDR) < 0.05 was considered significant). The data was obtained from the proteomics experiments and analysis presented in Koyuncu et al (2021)^6^.

| Protein | Total number of Ub-sites | Upregulated Ub-sites | Downregulated Ub-sites | Ub-sites with no changes | Total protein levels |
| --- | --- | --- | --- | --- | --- |
| IFB-2 | 3 | 0 | 1 | 2 | Increased |
| IFC-2 | 4 | 1 | 2 | 1 | No change |
| PAR-5 | 8 | 0 | 2 | 6 | No change |
| KIN-19 | 4 | 1 | 2 | 1 | No change |
| RHO-1 | 2 | 0 | 1 | 1 | Increased |
| DAF-21 | 11 | 3 | 4 | 4 | Increased |
| PAB-1 | 10 | 0 | 7 | 3 | Decreased |
| ATP-2 | 2 | 0 | 1 | 1 | No change |
| SCA-1 | 1 | 0 | 1 | 0 | No change |
| CPN-3 | 1 | 0 | 1 | 0 | Decreased |
| CCT-8 | 2 | 0 | 2 | 0 | No change |
| RACK-1 | 4 | 0 | 4 | 0 | No change |
| RPL-3 | 8 | 0 | 7 | 1 | No change |
| RPL-7 | 7 | 1 | 1 | 5 | Increased |
| RPL-8 | 2 | 0 | 1 | 1 | No change |
| RPL-10 | 4 | 0 | 3 | 1 | Decreased |
| RPL-17 | 8 | 0 | 5 | 3 | Decreased |
| RPL-23 | 2 | 0 | 1 | 1 | No change |
| RPL-31 | 2 | 0 | 2 | 0 | No change |
| RPS-0 | 1 | 0 | 1 | 0 | No change |
| RPS-5 | 0 | 0 | 0 | 0 | Decreased |
| RPS-11 | 3 | 0 | 2 | 1 | Increased |
| RPS-14 | 1 | 0 | 1 | 0 | Decreased |
| RPS-15 | 6 | 0 | 5 | 1 | No change |
| RPS-16 | 2 | 0 | 1 | 1 | No change |
| RPS-20 | 5 | 0 | 5 | 0 | Decreased |
| RPS-23 | 0 | 0 | 0 | 0 | Decreased |

We expanded our assessment to include other proteins reported to aggregate during aging in previous publications. In David et al., the authors established PAR-5, KIN-19, RHO-1, and DAF-21 as bona-fide examples of proteins that aggregate during aging^2^. Importantly, we could quantify all these insoluble proteins even in day 15-adult worms, and none of them were decreased at the total protein level (**Table 2**). Besides DAF-21, Reis-Rodrigues et al. identified other proteins that aggregate during aging, including PAB-1, ATP-2, SCA-1, CPN-3, CCT-8, RACK-1 and ribosomal subunits^3^. Our proteomics experiments successfully detected and quantified all these proteins in old worms (**Table 2**). Most of the proteins identified as aggregating during aging underwent ubiquitination changes at specific sites, primarily a loss of ubiquitination (**Table 2**). However, they also contained ubiquitinated sites that remained similar (**Table 2**).

In a recent preprint, Sui et al. performed limited proteolysis-coupled mass spectrometry (LiP-MS) to define changes in protein solubility across the proteome during aging and stress conditions in *C. elegans*^13^. After integrating their datasets with our ubiquitin proteomics data, the authors found that 91.8% of the age-dependent metastable proteins have differential ubiquitination during aging, while only 37.7% of them exhibit changes in phosphorylation^13^. This high correlation between metastable proteins and ubiquitination changes further supports that we assessed the global proteome, including aggregation-prone proteins

In our original publication^6^, we strengthened our conclusions regarding global changes in ubiquitination with western blot experiments. Using an antibody against ubiquitin (Sigma, 05-944, clone P4D1-A11), we observed a global decrease in ubiquitination levels from day 9 of adulthood^6^. In contrast, ubiquitination levels did not decrease in long-lived *eat-2* and *daf-2* mutant worms^6^. In addition to lysates from whole wild-type worms, we also found decreased ubiquitination levels during aging in isolated germlines, intestines, and heads, underscoring that ubiquitination changes occur across the entire organism. By combining various approaches including western blot analysis, we confirmed that loss of ubiquitination cannot be ascribed to a decrease in the total levels or half-life of ubiquitin itself. Given that a high proportion of DUBs were upregulated during aging in wild-type animals^6^, we examined whether elevated DUB activity contributes to ubiquitination changes. When we treated day-10 adult worms with broad-spectrum DUB inhibitor PR-619 for 4 hours before lysis, we observed a rescue of ubiquitination levels to those detected in day-1 adult worms^6^. In contrast, treating young worms with the DUB inhibitor for 4 hours did not induce a pronounced increase in their high ubiquitination levels^6^. Moreover, we found that knockdown of specific age-elevated DUBs (e.g., *csn-6, usp-5, H34C03*.*2*) ameliorate loss of ubiquitination during aging^6^. These data collectively reinforce our interpretation that there are changes in ubiquitination during aging and that this process is regulated.

The results discussed above (including ubiquitin proteomics and western blots) were presented in Figure 1 and associated extended Figures of our original publication^6^. In the other figures, we focused on age-dysregulated proteasome targets that accumulate during aging due to decreased ubiquitination and subsequent degradation by the proteasome. Particularly, we focused on two age-dysregulated proteasome targets: the IFB-2 intermediate filament and the EPS-8 regulator of RAC signaling. By proteomics and western blot, we confirmed that the protein levels of both IFB-2 and EPS-8 increase during aging^6^. Our microscopy and filter trap experiments demonstrated the aggregation of IFB-2 during aging, while a significant increase in its total levels was detected by proteomics and western blot. This example further supports that our experimental design does not overlook aggregated proteins. By employing a combination of approaches, we demonstrated that the accumulation of IFB-2 and EPS-8 during aging shortens lifespan, elucidating the molecular mechanisms underlying this process^6^. Importantly, knockdown of either IFB-2 or EPS-8 during adulthood was sufficient to extend longevity. Similar effects on lifespan regulation have also been reported in recent publications from other laboratories^14,15^. To confirm a direct link between loss of ubiquitination in IFB-2 and EPS-8 with longevity, we generated lysine to arginine mutants to block their ubiquitination. These Ub-less mutations were effective in suppressing degradation of endogenous IFB-2 and EPS-8 in young adults, accelerating their accumulation and subsequently shortening lifespan^6^. Together, these results provided functional evidence of a link between ubiquitination changes and aging, thereby validating the interpretation and conclusions of our original publication^6^.

**Fig. 1:**
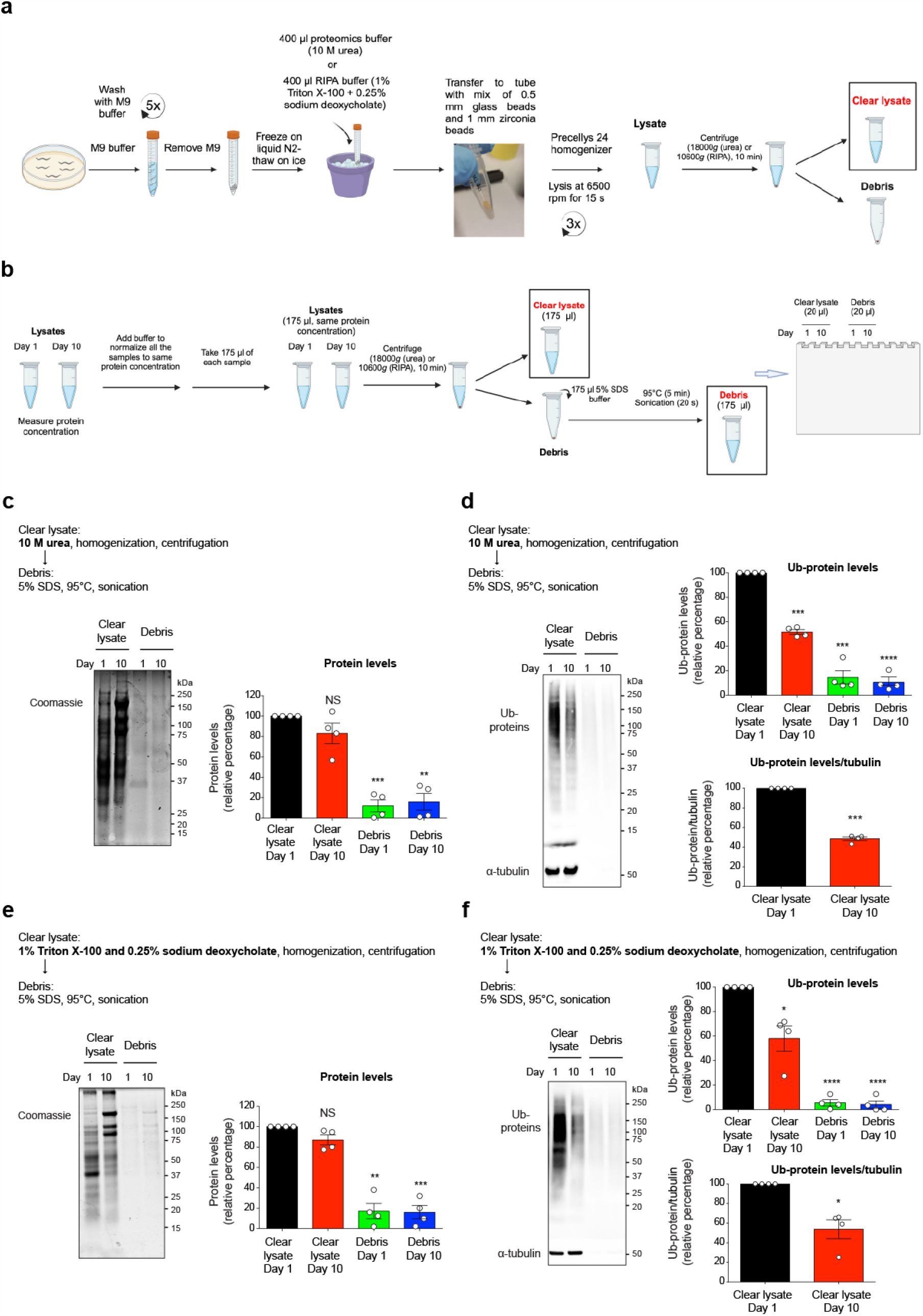
Our protein extraction protocols for proteomics and western blot experiments solubilize most of the total and ubiquitinated proteome. **a**, Scheme of worm lysis and protein extraction protocol used in our original publication^6^ and this manuscript. Approximately 2,000 living worms at different ages were collected using M9 buffer. After five washes with M9 buffer to eliminate bacteria, the worms were rapidly frozen with liquid nitrogen. Subsequent steps included thawing on ice, addition of lysis buffer (10 M urea buffer for proteomics; RIPA buffer with 0.25% sodium deoxycholate/1% Triton X-100 for western blot), and homogenization with a mix of 0.5 mm glass beads/1 mm zirconia beads using a Precellys 24 homogenizer (3 x 6,500 r.p.m. for 15 s). Both lysis buffers contained N-ethylmaleimide to inhibit deubiquitinating enzyme (DUB) activity. In our original publication, the resulting lysate was centrifuged for 10 min to eliminate worm debris (18,000g for proteomics; 10,600g for western blot). Clear lysates were collected, and protein concentration was determined for equal protein amount analysis. **b**, Scheme of the approach used to assess the proportion of total and ubiquitinated proteins remaining in the debris after centrifugation. Worm lysis and protein solubilization were carried out on day 1 and day 10-adult wild-type worms, following the protocol outlined in panel a. To ensure uniformity, all samples were normalized to the same protein concentration by adding lysis buffer. For each sample, 175 μl of homogenate underwent centrifugation, and the clear lysate was collected. The remaining proteins in the debris (175 μl) were extracted using 5% SDS RIPA buffer, followed by heating (95 °C, 5 min) and sonication. Subsequently, 20 μl of clear lysates and debris were loaded onto the same gel for comparison of total protein (Coomassie staining) and ubiquitinated protein levels (anti-ubiquitin antibody). **c**, Coomassie staining of clear lysates after protein extraction with 10 M urea and remaining debris solubilized with 5% SDS RIPA buffer, heating and sonication. Graph represents relative percentage values of protein levels to Day 1 Clear lysate (mean ± s.e.m., n= 4 independent experiments). **d**, Western blot with anti-ubiquitin antibody after protein extraction with 10 M urea and remaining debris solubilized with 5% SDS RIPA buffer, heating and sonication. α-tubulin is the loading control. The upper graph represents the relative percentage of ubiquitinated (Ub)-protein levels to Day 1 Clear lysate (mean ± s.e.m., n= 4 independent experiments). The lower graph represents the relative percentage (corrected for α-tubulin loading control) of Ub-protein levels in clear lysates to Day 1 (mean ± s.e.m., n= 4 independent experiments). **e**, Coomassie staining of clear lysates after protein extraction with 0.25% sodium deoxycholate/1% Triton X-100 RIPA buffer and remaining debris solubilized with 5% SDS RIPA buffer, heating and sonication. Graph represents relative percentage values of protein levels to Day 1 Clear lysate (mean ± s.e.m., n= 4 independent experiments). **f**, Western blot with anti-ubiquitin antibody after protein extraction with 0.25% sodium deoxycholate/1% Triton X-100 RIPA buffer and remaining debris solubilized with 5% SDS RIPA buffer, heating and sonication. The upper graph represents the relative percentage of Ub-protein levels to Day 1 Clear lysate (mean ± s.e.m., n= 4 independent experiments). The lower graph represents the relative percentage (corrected for α-tubulin loading control) of Ub-protein levels in clear lysates to Day 1 (mean ± s.e.m., n= 4 independent experiments). Statistical comparisons were made by two-tailed Student’s *t*-test for paired samples. *P* value: \**P* <0.05, \*\**P* <0.01, \*\*\**P* <0.001, \*\*\*\**P* <0.0001, NS, not significant (*P* >0.05).

### Analysis of total homogenates confirms loss of ubiquitination during aging

We appreciate the concerns raised by Daigle et al. and we have now performed additional tests to validate that the lysates analyzed in our initial publication are representative of global proteomes. To this end, we used the same lysis and protein extraction protocols as carried out in our original publication^6^ (please see **Fig. 1a** for detailed description). To clarify a concern from Daigle et al., we indeed removed dead worms before the lysis and we followed the same approach in the new tests. Briefly, approximately 2,000 worms at different ages were collected using M9 buffer and subjected to five washes with M9 buffer to eliminate bacteria. After removing the M9 buffer, the worms were rapidly frozen with liquid nitrogen. We thawed the worms on ice and then added the lysis buffer. For proteomics experiments, we used a buffer containing 10 M urea. For western blot analysis, we used RIPA buffer containing 0.25% sodium deoxycholate and 1% Triton X-100 as detergents. Protein ubiquitination is a reversible process, and this modification can be easily lost through the activity of DUBs as soon as cell lysis occurs^16^. In our original publication and the experiments presented here, we consistently included 25 mM N-ethylmaleimide in the lysis buffers to inhibit DUB activity and preserve ubiquitinated proteins as they existed in intact cells, following the standard in the field^16^.

The samples were transferred to tubes containing a mix of 0.5 mm glass beads and 1 mm zirconia beads for homogenization and protein extraction. To this end, we used Precellys 24 homogenizer (3 x 6,500 r.p.m. for 15 s) given its capacity to uniformly and efficiently homogenize different tissues and cells, ensuring high protein yield. In our original publication, the resulting lysate underwent centrifugation for 10 min to eliminate worm debris (10,600*g* for western blot, 18,000*g* for proteomics). Subsequently, we collected the resulting clear lysates and determined protein concentration. Equal amounts of total protein from these clear lysates were then subjected to analysis by either proteomics or western blot, as detailed in our original publication^6^.

To assess the remaining debris after centrifugation, we have now compared proportional amounts of clear lysates and debris (**Fig. 1b**). With this approach, we can determine the proportion of total and ubiquitinated proteins remaining in the debris compared to clear lysates. We conducted worm lysis and protein solubilization on day 1 and day 10-adult worms, following the above protocol (**Fig. 1a**). After lysis with Precellys 24 homogenizer, we measured the protein concentration in the resulting homogenates before centrifugation. We then normalized all samples to the same protein concentration by adding lysis buffer (**Fig. 1b**). For each sample, we centrifuged 175 μl of homogenate and collected the clear lysate. These lysates correspond to the samples analyzed in our original publication. The remaining proteins in the debris (175 μl) were extracted using 5% SDS RIPA buffer, followed by heating (95 °C, 5 min) and sonication, consistent with Daigle et al.’s method^1^ (**Fig. 1b**).

We loaded 20 μl of clear lysates and debris onto the same gel for comparison of total protein (Coomassie staining) and ubiquitinated protein levels (anti-ubiquitin antibody). In both day 1 and day 10-adult worms, we confirmed that 10 M urea solubilizes most of the proteome (**Fig. 1c**). Likewise, 10 M urea buffer solubilized most of the ubiquitinated proteome (**Fig. 1d**). Consequently, detecting ubiquitinated proteins in the debris was challenging compared to clear lysates (**Fig. 1d**). As reported in our original publication^6^, we observed a significant decrease (48.4%) in the levels of ubiquitinated proteins when comparing clear lysates from day 10-adult worms with day-1 adults (**Fig. 1d**). We obtained similar results (48.6% decrease in old worms) when the values were corrected for α-tubulin loading control in clear lysates (**Fig. 1d**).

Likewise, RIPA buffer containing 0.25% sodium deoxycholate and 1% Triton X-100 was also efficient at solubilizing most of the total and ubiquitinated proteome (**Fig. 1e-f**). Consistent with our original publication, the total amount of ubiquitinated proteins in clear lysates also decreased (46.2%) during aging when we used RIPA buffer containing 0.25% sodium deoxycholate and 1% Triton X-100 (**Fig. 1f**). However, the amount of ubiquitinated proteins remaining in the debris was almost undetectable when compared with clear lysates (**Fig. 1f**).

We then performed a second test to quantify age-related changes in total samples combining clear lysates and the remaining debris after protein extraction (**Fig. 2a**). Following the same worm lysis procedure as described above (**Fig. 1a**), we centrifuged 175 μl of homogenate after lysis with the Precellys 24 homogenizer. We separated the clear lysate, and the debris was solubilized in the same volume (175 μl) using 5% SDS RIPA buffer, followed by heating (95 °C, 5 min) and sonication. Then, we combined equal volumes of clear lysates and debris (60 μl of each) to obtain a total lysate. We measured protein concentration in the total lysates and analyzed equal protein amounts (20 μg) for each experimental condition (day 1 and day 10) (**Fig. 2a**). Importantly, we also observed a significant decline in ubiquitinated protein levels during aging when analyzing total lysates (**Fig. 2b,c**). This age-related decrease in ubiquitination, approximately 41%, remained consistent whether using 10 M urea or 0.25% sodium deoxycholate/1% Triton X-100 RIPA buffer (**Fig. 2b,c**). Similar to our findings in clear lysates^6^, treatment with the DUB inhibitor PR-619 in old worms for 4 hours before lysis restored ubiquitination levels (**Fig. 2d**). In conclusion, our data demonstrate that the samples analyzed in our original publication accurately represent changes in the whole proteome. This further validates the interpretation and conclusions drawn from the data presented in our original publication^6^, emphasizing alterations in the ubiquitinated proteome during aging.

**Fig. 2:**
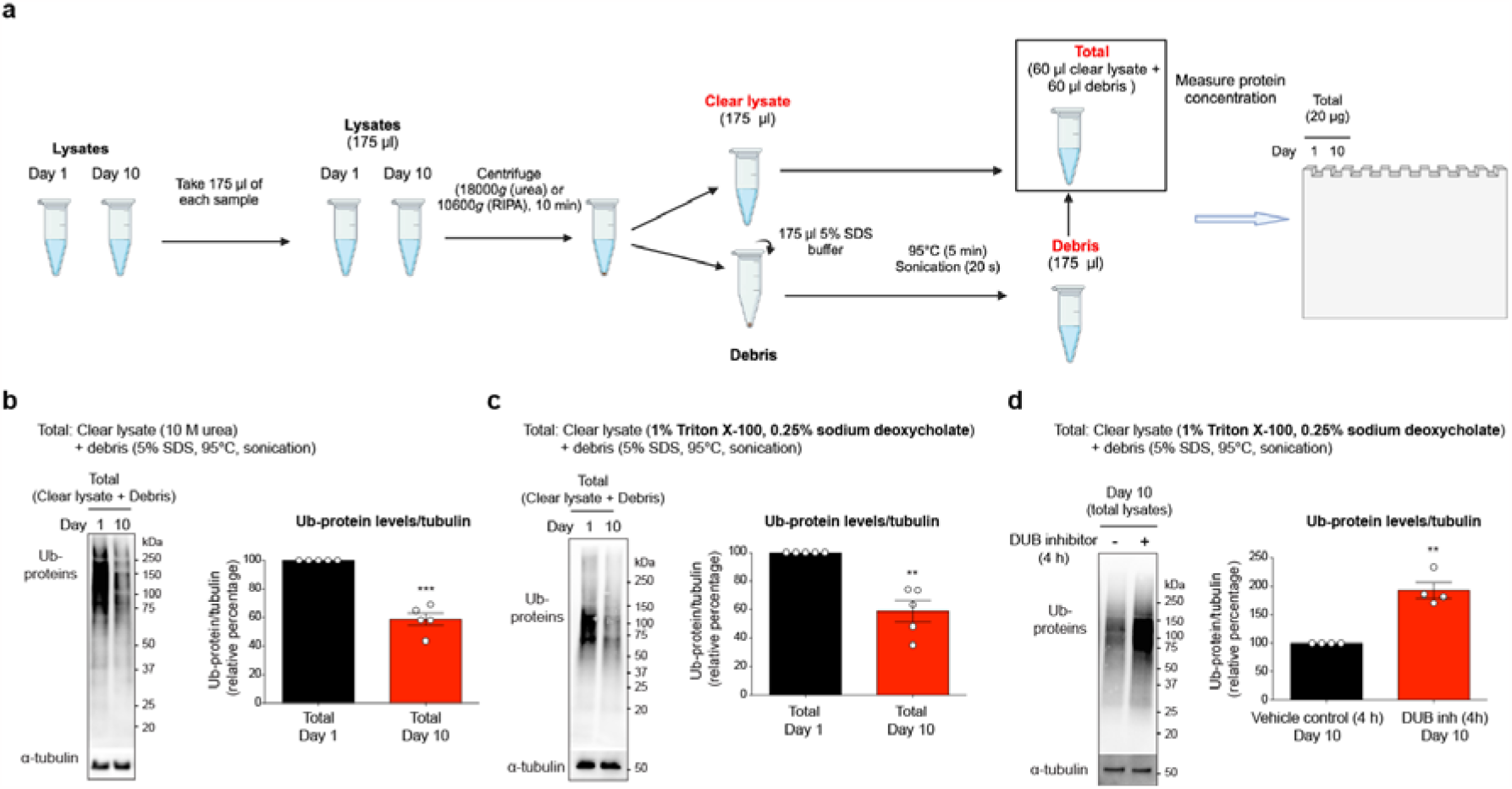
Analysis of total homogenates confirms loss of ubiquitination during aging. **a**, Scheme of the approach used to assess age-related changes in ubiquitinated proteins in total lysates. We centrifuged 175 μl of the homogenate obtained after lysis of wild-type worms using the Precellys 24 homogenizer. The clear lysate was separated, and the remaining debris was solubilized in the same volume (175 μl) using 5% SDS RIPA buffer, followed by heating (95 °C, 5 min) and sonication. Then, equal volumes of clear lysates and debris (60 μl each) were mixed to obtain a total lysate. We measured protein concentration in the total lysates. For each experimental condition (day 1 and day 10), we loaded 20 μg of total protein for further analysis. **b**, Western blot with anti-ubiquitin antibody of total homogenates combining clear lysates (10 M urea) and solubilized debris (5% SDS RIPA buffer, heating and sonication). α-tubulin is the loading control. Graph represents the relative percentage (corrected for α-tubulin loading control) of ubiquitinated (Ub)-protein levels to Day 1 (mean ± s.e.m., n= 5 independent experiments). **c**, Western blot with anti-ubiquitin antibody of total homogenates combining clear lysates (0.25% sodium deoxycholate/1% Triton X-100 RIPA buffer) and solubilized debris (5% SDS RIPA buffer, heating and sonication). Graph represents the relative percentage (corrected for α-tubulin loading control) of Ub-protein levels to Day 1 (mean ± s.e.m., n= 5 independent experiments). **d**, Immunoblot of Ub-proteins in total lysates from day 10-adult worms treated with 13.7 μg ml^−1^ PR-619 (broad-spectrum DUB inhibitor) or vehicle control (dimethyl sulfoxide, DMSO) for 4 h before lysis. Graph represents the relative percentage (corrected for α-tubulin loading control) of Ub-protein levels in total lysates to vehicle control (mean ± s.e.m., n= 4 independent experiments). Statistical comparisons were made by two-tailed Student’s *t*-test for paired samples. *P* value: \*\**P* <0.01, \*\*\**P* <0.001.

## Methods

### Synchronization of large *C. elegans* populations

In this study, wild-type *C. elegans* (N2) were cultured at 20□°C on standard Nematode Growth Medium (NGM) seeded with *E. coli* (OP50). To obtain synchronized populations of hermaphrodite worms for western blot analysis, we employed the bleaching technique. Young adult worms were collected by transferring random agar chunks from maintenance plates and allowed to grow until a sufficient number of young hermaphrodites were obtained for bleaching. Subsequently, the worms were treated with an alkaline hypochlorite solution (3 ml of bleach 4% (Fischer), 1.5 ml 5 N KOH, 5.5 ml dH_2_O) for 4 min to destroy adult tissues and obtain eggs. After multiple washes with M9 buffer, the eggs were maintained in M9 buffer without food overnight to enable hatching while preventing further development. Synchronized L1 larvae were randomly collected, raised and fed OP50 *E. coli* at 20□°C until late L4 larvae stage. Then, worms were transferred onto plates with OP50 *E. coli* covered with 100 μg ml^−1^ 5-fluoro-2′deoxyuridine (FUdR) to prevent progeny development. Adult worms were transferred onto fresh plates every five days.

### Worm lysis and protein extraction

Approximately 2,000 living worms at different ages were collected using M9 buffer and underwent five washes with M9 buffer to eliminate bacteria (**Fig. 1a**). After removal of the M9 buffer, the worms were rapidly frozen with liquid nitrogen. Then, we thawed the worms on ice and added the lysis buffer. Here we employed two distinct lysis buffers, consistent with our proteomics and western blot experiments in our initial publication^6^. The proteomics buffer consists of 10 M urea, 50 mM triethylammonium bicarbonate (TEAB) and 25 mM N-ethylmaleimide. The western blot buffer contains 50 mM Tris-HCl, pH 7.8, 150 mM NaCl, 1% Triton X-100, 0.25% sodium deoxycholate, 1 mM EDTA, 25 mM N-ethylmaleimide, 2 mM sodium orthovanadate, 1 mM PMSF and protease inhibitor cocktail (Roche). Subsequently, the samples were transferred to tubes containing a mixture of 0.5 mm glass beads and 1 mm zirconia beads for homogenization and protein extraction, using Precellys 24 homogenizer (3 x 6,500 r.p.m. for 15 s).

In our original publication, the resulting lysate underwent centrifugation for 10 min to eliminate worm debris (10,600*g* for western blot, 18,000*g* for proteomics). Clear lysates were then collected, and the protein concentration was determined for the analysis of equal protein amounts through proteomics or western blot. In this manuscript, we conducted additional tests to evaluate the proportion of total and ubiquitinated proteins remaining in the debris after centrifugation (**Fig. 1b**). Worm lysis and protein solubilization were conducted on day 1 and day 10-adult worms, following the above outlined protocol. After lysis with Precellys 24 homogenizer, we measured the protein concentration in the resulting homogenates with standard BCA protein assay (Thermo Scientific). We normalized all the samples to the same protein concentration by adding lysis buffer (**Fig. 1b**). For each sample, 175 μl of homogenate underwent centrifugation, and the clear lysate was collected. The remaining proteins in the debris (175 μl) were extracted using 5% SDS RIPA buffer, followed by heating (95 °C, 5 min) and sonication, consistent with Daigle et al.’s method^1^. Subsequently, 20 μl of clear lysates and debris were loaded onto the same 10% Bis-Tris gel for the comparison of total protein (Coomassie staining) and ubiquitinated protein levels (anti-ubiquitin antibody, Sigma, 05-944, clone P4D1-A11, 1:1,000, RRID: AB_441944). Additionally, the samples were analyzed with anti-α-tubulin (Sigma, T6199, 1:5,000, RRID: AB_477583) as a loading control.

In our second test, our goal was to evaluate age-related changes in ubiquitinated proteins in total samples combining clear lysates and the remaining debris after protein extraction (**Fig. 2a**). Following the same lysis procedure described previously (**Fig. 1a**), we centrifuged 175 μl of homogenate after lysis using the Precellys 24 homogenizer. The clear lysate was separated, and the resulting debris was solubilized in the same volume (175 μl) using 5% SDS RIPA buffer, followed by heating (95 °C, 5 min) and sonication. Then, we mixed equal volumes of clear lysates and debris (60 μl each) to obtain a total fraction. Subsequently, we determined the protein concentration in the total lysate using the BCA protein assay (Thermo Scientific). For each experimental condition (day 1 and day 10), we loaded 20 μg of protein from total lysates for further analysis.

### Coomassie Staining

The Bis-tris gel was covered with 20-30 ml of Coomassie solution. The gel was stained for 1 h at room temperature with gentle shaking. Then, the Coomassie solution was removed, and the gel was washed twice with deionized water for 5 min. Next, the gel was covered with destaining solution (9% Acetic Acid) and washed for 1 h with gentle shaking. Then, the destaining solution was removed, and the gel was covered with 50 ml of deionized water and washed overnight with gentle shaking.

## Acknowledgements

This work was supported by the Deutsche Forschungsgemeinschaft (DFG) (VI742/4-1 to D.V and Germany’s Excellence Strategy-CECAD, EXC 2030-390661388). The schemes of lysis protocols were generated using Biorender.

